# HIV-1 Nef and CycK:CDK13 antagonize SERINC5 for optimal viral infectivity

**DOI:** 10.1101/2021.05.24.445478

**Authors:** Qingqing Chai, Sunan Li, Morgan K. Collins, Rongrong Li, Iqbal Ahmad, Silas F. Johnson, Dylan A. Frabutt, Zhichang Yang, Xiaojing Shen, Liangliang Sun, Jian Hu, Judd F. Hultquist, B. Matija Peterlin, Yong-Hui Zheng

## Abstract

HIV-1 Nef antagonizes SERINC5 by redirecting this potent restriction factor to the endosomes and lysosomes for degradation. However, the precise mechanism remains unclear. Using affinity purification/mass spectrometry, we identified cyclin K and cyclin-dependent kinase 13 (CycK:CDK13) as a new Nef-associated kinase complex. CycK:CDK13 phosphorylates the serine at position 360 (S360) in SERINC5, which is required for Nef downregulation of SERINC5 from the cell surface and its counter activity of the SERINC5 antiviral activity. To understand the role of S360 phosphorylation, we created chimeric proteins between CD8 and SERINC5. Nef not only downregulates, but importantly, also binds to this chimera in a S360-dependent manner. Thus, S360 phosphorylation increases interactions between Nef and SERINC5 and initiates the destruction of SERINC5 by the endocytic machinery.

## Introduction

The accessory Negative Factor (Nef) protein is expressed by primate lentiviruses and functions to promote immune evasion, and subsequent viral fitness, by antagonizing a number of host cell surface proteins, such as CD4 and MHC class-I (MHC-I) (Kirchhoff, 2010). Recently, Nef was also found to antagonize three of the multi-pass transmembrane proteins representing the human SERine INCorporator (SERINC) family, whose five total members include SERINC1 through 5 (Inuzuka et al., 2005). SERINC5, and to a lesser degree, SERINC3, are specifically targeted by Nef in order to increase HIV-1 infectivity (Rosa et al., 2015; Usami et al., 2015). SERINC4 also has Nef-sensitive anti-HIV-1 activity, but it is rapidly turned over by host cell proteasomes (Qiu et al., 2020).

In a manner that is conserved across different species, SERINC5 is structurally composed of 10 transmembrane domains (TMDs), in an arrangement that forms five extracellular loops (ECLs), and four intracellular loops (ICLs) (Pye et al., 2020). In the absence of Nef, SERINC5 is incorporated into budding HIV-1 virions and subsequently inhibits viral entry into host cells by attacking HIV-1 Env trimers, in their open confirmation, thereby blocking the fusion pore formation between virions and target cells (Beitari et al., 2017; Chen et al., 2020; Sood et al., 2017; Zhang et al., 2019). The potency of SERINC5 antiretroviral activity has been validated in *SERINC5*-null mice (Timilsina et al., 2020).

HIV-1 Nef antagonizes the antiretroviral effect of SERINC5 by downregulating it from the cell surface, thereby preventing its virion incorporation, in a manner that is analogous to Nef-mediated CD4 downregulation (Rosa et al., 2015; Usami et al., 2015). The process is dependent on binding between the C-terminal dileucine motif ^160^ExxxLL^165^ of Nef and the adaptor protein 2 (AP-2) (Rosa et al., 2015) as well as an interaction of the N-terminal 32-39 amino acids (aa) of Nef with SERINC5 (Ananth et al., 2019). Amino acids 32-39 of Nef are located in a multifunctional hydrophobic pocket that is also required for binding to the cytoplasmic tails of CD4 and MHC-I (Kwon et al., 2020). The assembly of SERINC5 and CD4 with the Nef/AP2 complex leads to their endocytosis and subsequent lysosomal mediated degradation (daSilva et al., 2009; Shi et al., 2018).

The importance of Nef mediated antagonism of SERINC5 is illustrated by both its conservation across primate lentiviruses and the fact that the global spread of primate lentiviruses is directly correlated to the potency of Nef-mediated SERINC5 antagonism (Heigele et al., 2016). SERINC5 is also counteracted by murine leukemia virus (MLV) glycoGag and by the equine infectious anemia virus (EIAV) S2 protein (Ahi et al., 2016; Chande et al., 2016; Rosa et al., 2015). Although glycoGag and S2 do not share any homology with Nef, they also target SERINC5 to endosomes and lysosomes for degradation (Ahmad et al., 2019; Li et al., 2019). Finally, the clinical relevance of SERINC5 antagonism is underscored by the observation that *Nef* genes isolated from HIV-1 controllers antagonize SERINC5 poorly when compared to those from progressors (Jin et al., 2019). Thus, SERINC5 is an important restriction factor whose antiretroviral activity has been demonstrated *in vitro* and *in vivo*.

Cyclin-dependent kinases (CDKs) belong to a family of serine/threonine kinases whose activities are regulated by cyclin (Cyc) subunits (Lim and Kaldis, 2013). The human genome encodes 21 CDKs and 29 Cycs, the functions of which can be separated into two major categories. Whereas CDK1/2, CDK4, and CDK6 regulate the cell cycle, CDK7-9, CDK11, and CDK12/13 play essential roles in transcription (Fan et al., 2020). CDK12, in conjunction with the regulatory subunit CycK (Bartkowiak et al., 2010; Blazek et al., 2011), phosphorylate the C-terminal domain in RNA polymerase II (Bosken et al., 2014; Greifenberg et al., 2016) and consequently regulates genome stability and RNA processing. Although CDK13 mediated phosphorylation proceeds in a manner similar to CDK12 via CycK, the precise cellular function of CDK13 remains largely unknown (Greenleaf, 2019). In this study, we demonstrate that CycK:CDK13 phosphorylates SERINC5 and this phosphorylation is required for Nef antagonism of SERINC5.

## Results

### Identification of CycK:CDK13 as a Nef-associated kinase via mass spectrometry

Initially, FLAG-tagged SERINC5 protein was expressed alone in HEK293T cells and purified by an anti-FLAG affinity column. When the eluted proteins were analyzed by SDS-PAGE, purified monomeric SERINC5 with a molecular mass ∼45 kDa was detected via Coomassie Brilliant Blue staining (**Fig.1A**). Next, SERINC5 was purified again after its co-expression with wild-type (WT) and Nef-deficient (ΔNef) HIV-1 proviral vectors. To avoid the degradation of SERINC5 by Nef, the amounts of proviral vectors were minimized. The monomeric SERINC5 was purified again, but notably, a ∼170-kDa protein was co-purified with it only in the presence of Nef (**Fig.1B,** lane 3, labelled with *). To identify this large protein, purified proteins were analyzed by liquid chromatography tandem mass spectrometry (LC-MS/MS). CDK13 was one of six large proteins identified that ranged in size from 150 to 200 kDa (**Fig.1C**). Because CDK13 interacts with CycK, and importantly, CycK is a well-characterized Nef-binding protein (Khan and Mitra, 2011), CycK:CDK13 was selected for further study.

**Figure 1.**
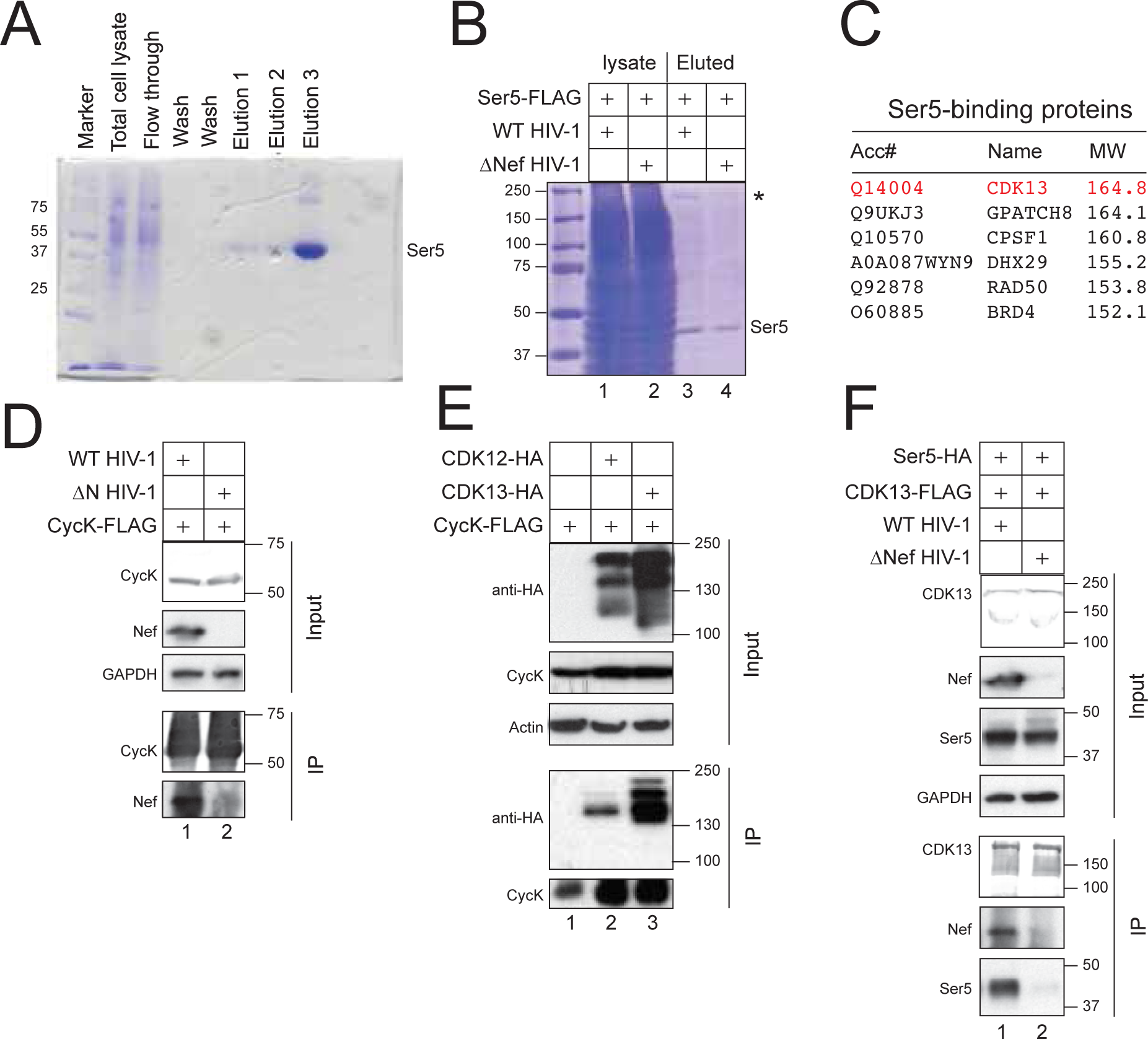
Identification of CycK:CDK13 as a Nef-associated kinase via mass spectrometry. A) FLAG-tagged SERINC5 protein was expressed in HEK293T cells and purified by anti-FLAG M2 affinity chromatography. After SDS-PAGE, proteins from total cell lysate and three eluted fractions were analyzed after being stained with Coomassie Brilliant Blue. Monomeric SERINC5 proteins are indicated. B) FLAG-tagged SERINC5 protein was expressed with WT and ΔNef HIV-1 proviral vectors in HEK293T cells and purified and analyzed similarly as in A). Monomeric SERINC5 and a protein of ∼170 kDa (*) are indicated. C) Purified proteins from B) were further analyzed by liquid chromatography/mass spectrometry (LC/MS). Control experiments were conducted using beads that were not conjugated with any antibodies. Six proteins with molecular weights 150-200 kDa that are not found in the control experiments are listed. D) FLAG-tagged CycK protein was expressed with WT and ΔNef HIV-1 proviral vectors in HEK293T cells. Proteins were immunoprecipitated with anti-FLAG antibodies and analyzed by western blotting (WB). CycK was detected by anti-FLAG antibodies, and Nef and GAPDH by specific antibodies. IP, immunoprecipitation; Input, cell lysate. E) FLAG-tagged CycK protein was expressed with HA-tagged CDK12 or CDK13 proteins in HEK293T cells. Proteins were immunoprecipitated and analyzed as in D). CDK12 and CDK13 were detected by anti-HA antibodies. F) FLAG-tagged CDK13 protein was expressed with HA-tagged SERINC5 protein in the presence of WT and ΔNef HIV-1 proviral vectors in HEK293T cells. Proteins were immunoprecipitated and analyzed as in D) and E).

First, we co-expressed FLAG-tagged CycK protein with WT or ΔNef HIV-1 proviral vectors. Proteins were pulled down by anti-FLAG antibodies and analyzed by western blotting (WB). CycK specifically pulled down Nef (**Fig.1D**, lane 1). The same CycK was also co-expressed with HA-tagged CDK12 or CDK13 proteins. Two major CDK12 and CDK13 species were detected at 130-200 kDa. CycK pulled down the low molecular weight CDK12 and both CDK13 species (**Fig.1E**, lanes 2-3). These results confirmed Nef-CycK, CycK:CDK12, and CycK:CDK13 interactions as reported previously.

Second, we expressed FLAG-tagged CDK13 with HA-tagged SERINC5 proteins in the presence of reduced amounts of WT and ΔNef HIV-1 proviral vectors. When proteins were pulled down again by anti-FLAG antibodies, CDK13 associated with SERINC5 only in the presence of Nef (**Fig.1F,** lane 1). Thus, Nef is an adaptor that bridges SERINC5 with CycK:CDK13.

### CycK:CDK13 is required for Nef downregulation of SERINC5

We examined how CycK:CDK13 affects Nef downregulation of SERINC5 by silencing the endogenous gene expression of CycK (encoded by CCNK). Because CycK is essential for cell survival (Dai et al., 2012), we used CRISPR interference (CRISPRi). Catalytically dead Cas9 (dCas9) and *CCNK*- specific guide RNA (gRNA) were co-expressed in 293T cells, and their silencing of the endogenous CycK was assessed by WB. Unlike the ectopic CycK protein that was detected at ∼55 kDa, endogenous CycK protein was detected at ∼55 kDa and ∼75 kDa (**Fig.S1A**). Both CycK species were reduced by *CCNK*-CRISPRi in a dose-dependent manner (**Fig.S1A**).

*CCNK*-knockdown (KD) was reported to stall cell-cycle in the G1 phase (Lei et al., 2018). Indeed, when dCas9/*CCNK*-gRNA were co-expressed in human CD4^+^ Jurkat T cells, the fraction of cells in G2 were decreased in a dose-dependent manner (**Fig.S1B**). Importantly, under such *CCNK*-KD condition, Nef was unable to decrease SERINC5 protein expression in 293T cells (**Fig.2A,** upper panel, lanes 1, 3). We reported that Nef also decreases SERINC5 protein expression in Jurkat cells (Shi et al., 2018). When the endogenous CycK was similarly targeted by *CCNK*- CRISPRi in Jurkat cells, the Nef effect also diminished (**Fig.2A,** lower panel). Measurements of SERINC5 expression on the cell surface produced similar results. Nef effectively decreased both the number of SERINC5-positive cells (**Fig.2B**) and the mean fluorescence intensity (MFI) of these positive cell populations (**Fig.2E**) Nef also downregulates CD4 from the cell surface (Shi et al., 2018), but it does not target SERINC2 (Dai et al., 2018). However, *CCNK*-KD had no effect on Nef downregulation of CD4 expression, and neither did it affect SERINC2 expression (**Fig.S1C**, lanes 1, 3, 5, 7). Thus, the contribution of CycK is highly specific for the effect of Nef on SERINC5.

**Figure 2.**
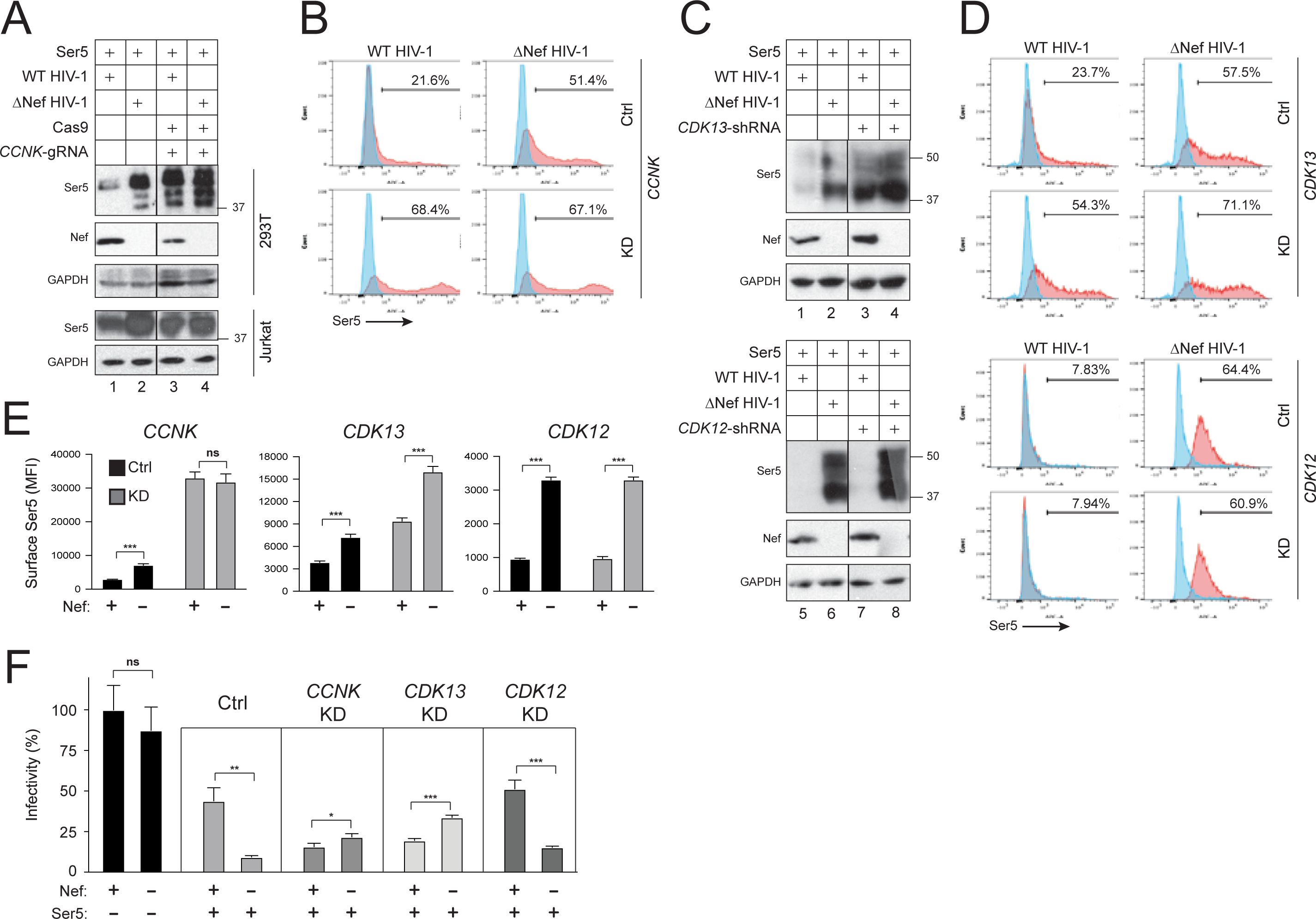
CycK:CDK13 is required for Nef downregulation of SERINC5. A) SERINC5 was expressed with Cas9/*CCNK*-gRNA expression vectors in the presence of WT and ΔNef HIV-1 proviral vectors in HEK293T (upper gels), or *SERINC3/5* knockout Jurkat-TAg cells (lower gels). Proteins were analyzed by WB. SERINC5 was detected by anti-FLAG antibodies. B) SERINC5 with an internal HA-tag (Ser5-iHA) was expressed with *CCNK-*shRNA expression vector in the presence of WT and ΔNef HIV-1 proviral vectors in HEK293T cells. SERINC5 expression on the cell surface was analyzed by flow cytometry using fluorescent anti-HA antibodies. Results are presented as histograms and SERINC5-positive cell populations are indicated (%). C) SERINC5 was expressed with *CDK12*- or *CDK13*-shRNA expression vectors in the presence of WT and ΔNef HIV-1 proviral vectors in HEK293T, and protein expression was analyzed by WB. SERINC5 was detected by anti-FLAG antibodies. D) Ser5-iHA was expressed with *CDK12*- or *CDK13*-shRNA expression vectors as in B) and analyzed similarly. E) Mean fluorescence intensity (MFI) values for the SERINC5-positive cell populations in B) and D) were statistically analyzed. F) SERINC5 was expressed with WT and ΔNef HIV-1 proviral vectors in the presence of indicated shRNA or control (Ctrl) expression vectors in 293T cells. After virions were collected and quantified by p24^Gag^ ELISA, TZM-bI cells were infected with an equal amount of virions and viral infectivity was analyzed by measuring the intracellular firefly luciferase activity after 48 hours of infection. Infectivity is presented as a relative value, with the WT HIV-1 infectivity in absence of SERINC5 set to 100%. Error bars in E) and F) represent the SEM from three independent experiments. Statistical analysis: *p<0.05, **p<0.01, ***p<0.001, ns, not significant (p>0.05).

Next, we examined the endogenous CDK13 activity using CDK12 as a control. Like CycK, both CDK12 and CDK13 are essential for cell survival, so we used previously reported short hairpin RNAs (shRNAs) to knock down their expression (Dai et al., 2012). Endogenous CDK12 was detected as a single species at ∼180 kDa, and endogenous CDK13 was detected as multiple species from ∼150 to 200 kDa (**Fig.S2**). *CDK12*-shRNA and *CDK13*-shRNA effectively reduced the expression of their target proteins (**Fig.S2**). Notably, Nef no longer decreased the expression of SERINC5 in the presence of *CDK13*-KD (**Fig.2C**, lanes 1-4). In contrast, this effect of Nef was not affected by *CDK12*-KD (**Fig.2C**, lanes 5-8). In addition, *CDK13*-KD disrupted the downregulation of SERINC5 by Nef from the cell surface, whereas *CDK12*-KD did not (**Fig.2D, Fig.2E**).

Furthermore, we determined how *CCNK*-KD, *CDK12*-KD, and *CDK13*-KD affect Nef counter activity of the SERINC5 anti-HIV-1 activity using a viral infectivity assay. SERINC5 strongly reduced the ΔNef HIV-1 infectivity, which was counteracted by Nef (**Fig.2F**). This Nef counter activity was disrupted by *CCNK*- and *CDK*13-KD, but not by *CDK12*-KD (**Fig.2F**). Taken together, the KD of both endogenous CycK and CDK13 support the conclusion that CycK:CDK13 is required for the downregulation of SERINC5 by Nef.

### The serine 360 residue in SERINC5 is critical for Nef antagonism of SERINC5

CDKs preferentially phosphorylate serine or threonine residues in the motif [(S/T)Px(R/K)], in which only the (S/T)P residues are invariant (Lim and Kaldis, 2013). In SERINC5, the only two (S/T)P motifs present in the protein are found at AA positions ^249^SP^250^ and ^360^SP^361^. Due to its presence in TMD6, S249 is unlikely to be accessible for phosphorylation by CycK:CDK13 (**Fig.3A**). In contrast, the presence of S360 in ICL4 suggests that this residue is a likely target for phosphorylation (**Fig.3A**). In order to evaluate the importance of S249 and S360 on Nef downregulation of SERINC5, we independently substituted both residues to alanine and tested their respective response to Nef. When WT, S249A, or S360A SERINC5 proteins were co-expressed with WT or ΔNef HIV-1 proviral vectors in HEK293T cells, Nef selectively decreased the expression of WT and S249A SERINC5, but not S360A SERINC5 at steady-state levels (**Fig.3B,** lanes 2, 4, 6).

**Figure 3.**
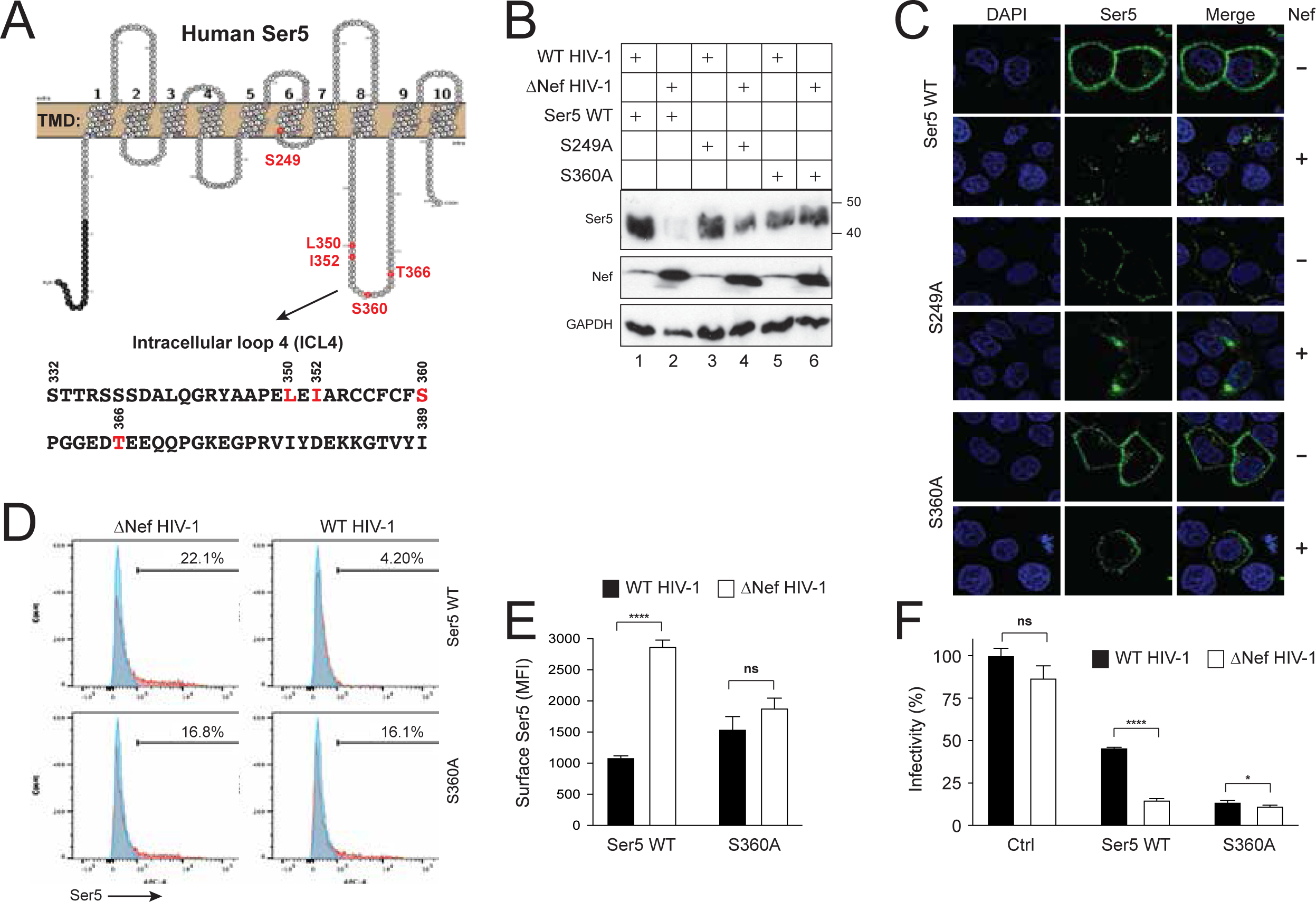
The serine 360 residue in SERINC5 is critical for Nef antagonism of SERINC5. **A)** SERINC5 transmembrane topology and its ICL4 amino acid sequence are presented. Residues selected for mutagenesis are indicated in red. **B)** WT, S249A, and S360A SERINC5 proteins were expressed with WT or ΔNef HIV-1 proviral vectors in HEK293T cells. SERINC5 proteins were detected by WB with anti- FLAG antibodies. **C)** WT, S249A, and S360A SERINC5 proteins with an internal FLAG-Tag were expressed with WT or ΔNef HIV-1 proviral vectors in HeLa cells. Cells were stained with fluorescent anti-FLAG antibodies, and the antibody uptake was determined by confocal microscopy at 37°C. **D)** WT and S360A SERINC5 proteins with internal HA-tags were expressed with WT or ΔNef HIV-1 proviral vectors in HEK293T cells and analyzed by flow cytometry. **E)** MFI values for the SERINC5-positive cell populations in D) were statistically analyzed. **F)** The anti-HIV-1 activity of WT and S360A SERINC5 proteins were compared and presented similarly as before. Error bars represent the SEM from three independent experiments. Statistical analysis: *p<0.05, ****p<0.0001, ns p>0.05.

Three more experiments were conducted to confirm the effect of the S360A substitution. First, the WT, S249A, or S360A SERINC5 proteins were co-expressed with WT and ΔNef HIV- 1 proviral vectors in HeLa cells, and SERINC5 endocytosis was determined by a confocal microscopy-based antibody uptake assay (Shi et al., 2018). WT and S249A SERINC5 were endocytosed in the presence of Nef, whereas S360A SERINC5 was not (**Fig.3C,** top 12 panels versus bottom 6 panels). Second, Nef downregulation of WT and S360A SERINC5 from the cell surface were measured by flow cytometry. Although WT SERINC5 was effectively downregulated in the presence of Nef, the S360A SERINC5 protein was not (**Fig.3D, Fig.3E**). Third, the antiviral activity of SERINC5 was determined similarly as in Fig.2F. Whereas WT SERINC5 strongly reduced ΔNef HIV-1 infectivity, S360A SERINC5 strongly inhibited both WT and ΔNef HIV-1 infectivity (**Fig.3F**). These results demonstrate that the S360 residue is required for Nef-mediated downregulation and its counter activity of SERINC5.

### A phosphomimetic S360D substitution reduces the expression of SERINC5

It was previously reported that two hydrophobic residues in the ICL4 region of SERINC5 (L350, I352) also determine sensitivity of the protein to Nef (Dai et al., 2018). Therefore, we substituted both L350 and I352 residues to alanine in order to generate L350A/I352A SERINC5 and subsequently tested its sensitivity to Nef, compared to WT and S360A SERINC5. When the WT, S360A, and L350A/I352A SERINC5 proteins were expressed alone in Jurkat cells, they all had similar levels of cell surface expression (**Fig.4A, Fig.4B**). When they were individually expressed with WT and ΔNef HIV-1 proviral vectors, WT SERINC5 was sensitive, whereas the S360A and L350A/I352A SERINC5 proteins were resistant to Nef-induced downregulation from the cell surface (**Fig.4C, Fig.4D**). These results were also confirmed in 293T cells (**Fig.S3**), indicating that their resistance Nef is not cell type-dependent.

**Figure 4.**
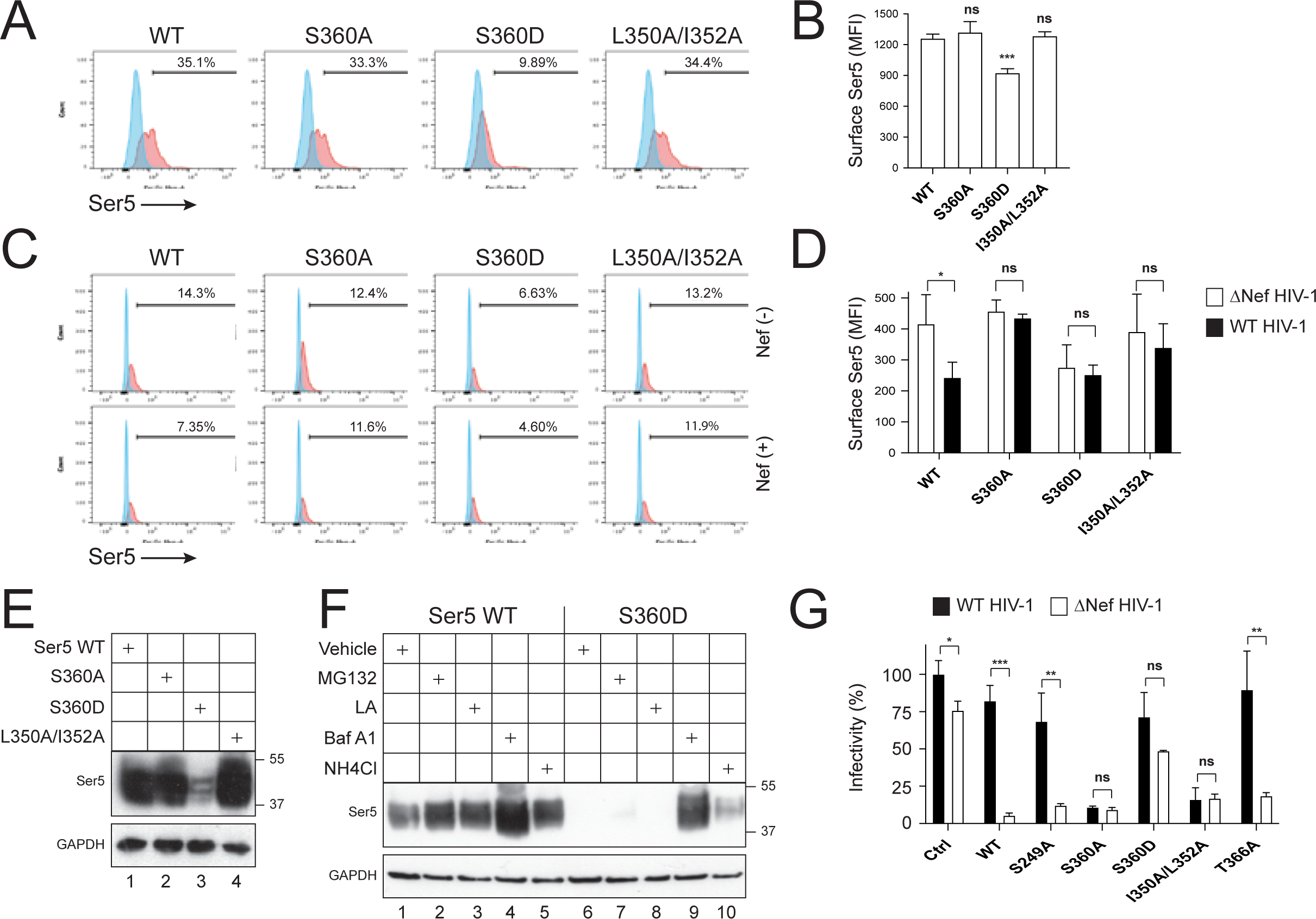
A phosphomimetic S360D substitution reduces the expression of SERINC5. **A)** WT, S360A, S360D, and L350A/I352A SERINC5 proteins with internal HA-tags were expressed in *SERINC3/5* knockout Jurkat-TAg cells. SERINC5 expression on the cell surface was determined by flow cytometry. **B)** MFI values for the SERINC5-positive cell populations in A) were statistically analyzed. **C)** WT, S360A, S360D, and L350A/I352A SERINC5 proteins with internal HA-tags were expressed with WT or ΔNef HIV-1 proviral vectors in *SERINC3/5* knockout Jurkat-TAg cells. SERINC5 expression on the cell surface was determined by flow cytometry. **D)** MFI values for the SERINC5-positive cell populations in C) were statistically analyzed. **E)** Indicated SERINC5 proteins with an internal HA-tag were expressed in HEK293T cells and their expressions were determined by WB using anti-HA antibodies. **F)** WT and S360A SERINC5 proteins were expressed in HEK293T cells. Cells were treated with MG132, lactacystin (LA), bafilomycin A1 (Baf A1), or NH4Cl, and SERINC5 protein expression was determined by WB. **G)** The anti-HIV-1 activity of indicated SERINC5 proteins were compared and presented similarly as we did before. Error bars in B), D), and G) represent the SEM from three independent experiments. Statistical analysis: *p<0.05, **p<0.01, ***p<0.001, ns p>0.05.

Next, we continued to examine the important role of S360 in SERINC5 by creating a phosphomimetic protein which contained a serine-to-aspartic acid (S360D) substitution. When expressed in Jurkat cells, regardless of Nef presence, the S360D SERINC5 cell surface expression was lower than the WT, S360A, or L350A/I352A SERINC5 proteins (**Fig.4A-Fig.4D**). When we compared their expression at steady-state by WB in HEK293T cells, we also observed a dramatic decrease in the expression of S360D SERINC5 (**Fig.4E**, lane 3). To determine if the decreased levels of S360D SERINC5 were a result of protein degradation, cells were treated with proteasomal inhibitors (MG132, lactacystin) and lysosomal inhibitors (Bafilomycin A1, NH4Cl). The expression of S360D SERINC5 protein was increased strongly by bafilomycin A1 and mildly by NH4Cl, whereas MG132 and lactacystin barely had any effect (**Fig.4F**, lanes 6-10). On the contrary, although WT SERINC5 expression levels were increased in the presence of all four inhibitors, their relative increase was less than that of S360D SERINC5 (**Fig.4F**, lanes 1-5). Thus, the phophomimietic S360D SERINC5 is selectively targeted to lysosomes for degradation.

Previously, the T366 residue in ICL4 of SERINC5 was shown to be phosphorylated by casein kinase II *in vitro* (Stoneham et al., 2020). Therefore, we constructed a T366A SERINC5 to further complement our panel of SERINC5 proteins. Expression of the WT, S249A, or T366A SERINC5 proteins strongly reduced HIV-1 infectivity only in the absence of Nef, whereas S360A and L350A/I352A SERINC5 proteins did so independently of Nef (**Fig.4G**). The marginal antiviral activity S360D SERINC5 was due to its relative low levels of expression. Collectively, these results confirm that residues S360, L350, and I352 determine the sensitivity of SERINC5 to Nef-mediated downregulation and demonstrate that a negative charge at position 360 destabilizes SERINC5, due to its phosphomimetic properties. As such, they point to an important role of S360 phosphorylation in Nef antagonism of SERINC5.

### CycK:CDK13 phosphorylates the serine 360 residue in SERINC5

In the CDK13 kinase domain, D837 acts as proton acceptors for phosphate transfer to the substrate, and D855 binds to two Mg^2+^ ions for coordinating the phosphate donor. Simultaneous disruption of these two residues by introducing D837A/D855N substitutions completely abrogates the CDK13 kinase activity (Greifenberg et al., 2016). To determine whether the S360 residue of SERINC5 is phosphorylated by CycK:CDK13, we generated D837A and D837A/D855N substitutions in CDK13 and tested their activities on Nef downregulation of SERINC5. In addition, we generated similar D858A and D858A/D876N substitutions in CDK12 and tested their activities. Both D837A CDK13 and D858A CDK12 proteins had not effect on Nef-mediated downregulation of SERINC5 (**Fig.5A**, lanes 1-2, 5-6). However, the D837A/D855N CDK13 protein disrupted Nef activity, whereas the D858A/D876N CDK12 did not, suggesting that CDK13 kinase activity is critical for Nef-mediated SERINC5 downregulation (**Fig.5A**, lanes 3-4, 7-8). To confirm this, we treated cells with a CDK12/CDK13 covalent inhibitor THZ531 (Zhang et al., 2016). We found the activity of Nef was also disrupted by this inhibitor (**Fig.5B**, lane 3). Taken together, these results demonstrate that CDK13 kinase activity is required for the downregulation of SERINC5 by Nef.

**Figure 5.**
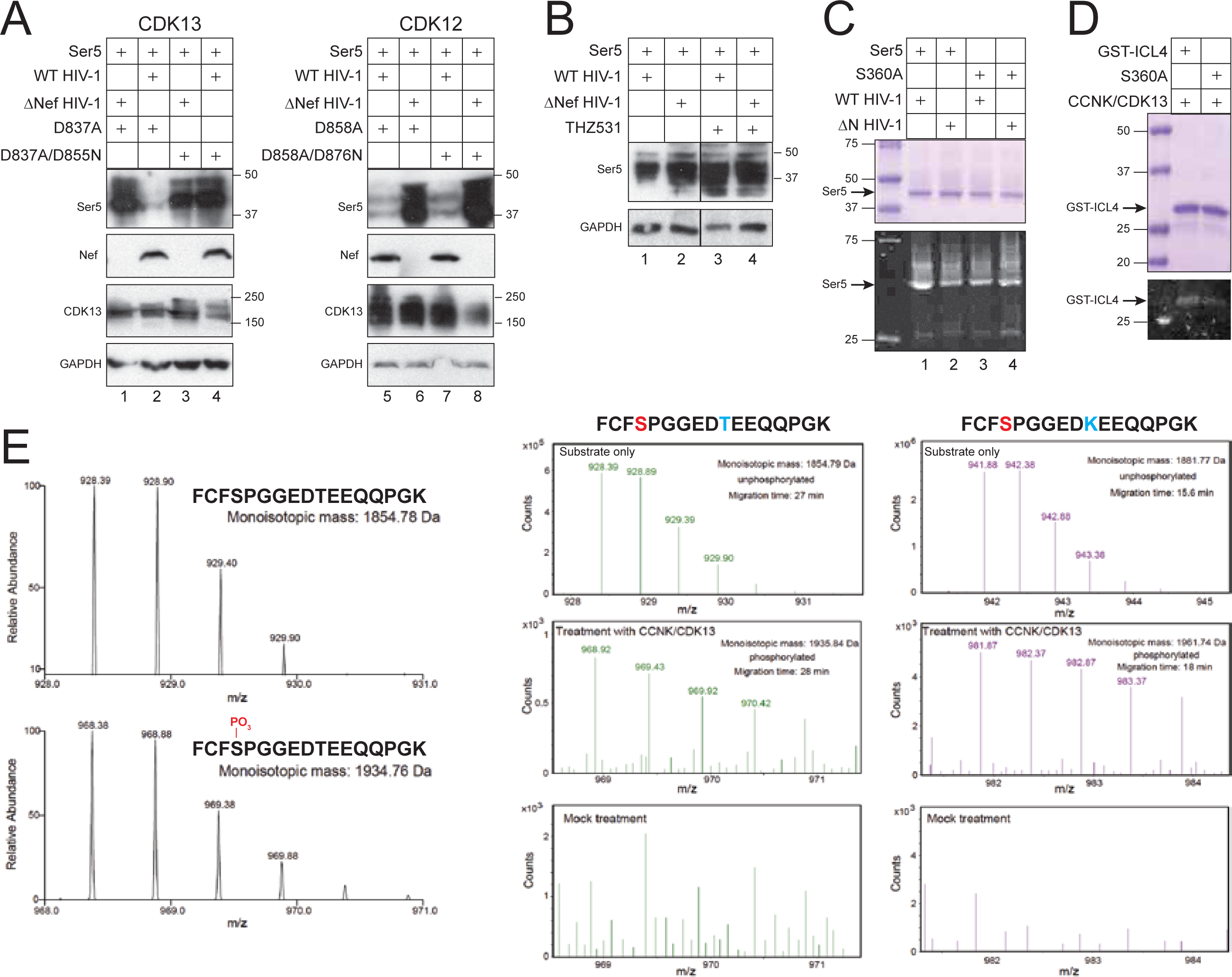
CycK:CDK13 phosphorylates the serine 360 residue in SERINC5. **A)** SERINC5 was expressed with indicated CDK13 or CDK12 proteins in the presence of WT or ΔNef HIV-1 proviral vectors in HEK293T cells. SERINC5, CDK13, and CDK12 proteins were detected by WB using anti-FLAG antibodies. **B)** SERINC5 was expressed with WT or ΔNef HIV-1 proviral vectors in HEK293T cells and treated with THZ531 at 100 μM. SERINC5 expression was analyzed by WB using anti- FLAG antibodies. **C)** WT and S360A SERINC5 proteins were expressed with WT or ΔNef HIV-1 proviral vectors in HEK293T cells and purified similarly as before. After SDS-PAGE, proteins were analyzed after being stained with Coomassie Brilliant Blue or Phos-Tag. **D)** WT and S360A GST-ICL4 recombinant proteins were expressed in *E.coli* and purified by glutathione resin. *In vitro* kinase assay was conducted by incubating these proteins with affinity purified CycK:CDK13. After SDS-PAGE, proteins were analyzed after being stained with Coomassie Brilliant Blue or Phos-Tag. **E)** Peptide ^357^FCFSPGGEDTEEQQPGK^373^, its S360 phosphorylated version, and its T366K substitution version were synthesized. They were analyzed by high-resolution MS via direct infusion, or analyzed by CZE-MS after treatment with affinity purified CycK:CDK13.

To determine whether SERINC5 is phosphorylated at S360 in cells, we purified WT and S360A SERINC5 proteins from HEK293T cells in the presence of HIV-1 proviral vectors as described previously (**Fig.5C**, upper gel). When their phosphorylation status was analyzed by Phos-Tag staining, WT and S360A SERINC5 proteins showed similar levels of phosphorylation in the absence of Nef (**Fig.5C**, lower gel, lanes 2, 4). However, in the presence of Nef, the WT SERINC5 protein phosphorylation was increased relative to S360A SERINC5 (**Fig.5C**, lower gel, lanes 1, 3).

To confirm that S360 is phosphorylated by CycK:CDK13, we conducted an *in vitro* kinase assay (Greifenberg et al., 2016). The ICL4 regions of both WT and S360A SERINC5 were independently fused to the glutathione S-transferase (GST) tag. WT and S360A GST-ICL4 recombinant proteins were expressed in *E.coli* and purified by glutathione-Sepharose beads (**Fig.5D**). In addition, the CycK:CDK13 complex was directly purified from HEK293T cells by anti-FLAG affinity chromatography. When purified proteins were incubated with CycK:CDK13 and analyzed by Phos-Tag staining, the WT GST-ICL4 protein showed a relatively stronger phosphorylation signal than the S360A GST-ICL4 protein (**Fig.5D**).

To confirm this result, we conducted another *in vitro* kinase assay, using ICL4-derived 17- aa peptides as substrates, and measured the phosphorylation by high-resolution MS via direct infusion. We compared the unphosphorylated WT peptide (^357^FCFSPGGEDTEEQQPGK^373^) with the S360 phosphorylated version (**Fig.5E,** left two panels) and, as expected, found that the phosphorylated peptide had an 80-Da higher mass than the WT peptide (1934.76 Da *vs.* 1854.78 Da).

We then used WT ICL4-derived peptide and a control peptide containing a T366K substitution (^357^FCFSPGGED<u>K</u>EEQQPGK^373^) to conduct the kinase assay and analyzed the products by capillary zone electrophoresis (CZE)-MS. In the WT sample, a peptide with an ∼81- Da higher mass, compared to the mock treatment control, was observed (**Fig.5E**, middle panels). This result was most likely due to the phosphorylation (+80 Da) and deamination (+1 Da) of the peptide. Moreover, the phosphorylated peptide migrated slower than the unphosphorylated control during CZE separation due to the addition of one negative charge on the peptide (28 *vs.* 27 min), a result which is consistent with previous phosphoproteomic data of CZE-MS (Chen et al., 2019). In the T366K sample, a phosphorylated version of the peptide was also detected (**Fig.5E**, right panels). Similarly, this peptide had an 80-Da higher mass than the unphosphorylated one (1961.74 Da *vs.* 1881.77 Da) and migrated significantly slower than the unphosphorylated control (18 min vs. 15.6 min). Due to the fact that the T366K substitution eliminates that residue as a potential phosphorylation target, this result suggests that the phosphorylation of the T366K peptide also occurred at the S360 residue. Collectively, these results demonstrate that CycK:CDK13 phosphorylates S360 in SERINC5.

### Nef downregulates CD8-ICL4 fusion protein

CD8 is a type I membrane protein with a signal peptide (SP), an N-terminal extracellular domain (ECD), a TMD, and a short C-terminal cytoplasmic tail (CT). CD8 exists as an αα homodimer or an αβ heterodimer, and Nef selectively downregulates CD8β by interacting with its CT (Stove et al., 2005). Previously, we created CD8α- Nef fusion protein to study Nef-mediated internalization of cell surface molecules (Mandic et al., 2001). To further characterize the role of phosphorylation on Nef-mediated downregulation of SERINC5, we generated a WT CD8α-ICL4 fusion protein by both swapping the CT of CD8α with the ICL4 of SERINC5 and inserting an HA-Tag immediately downstream of SP (**Fig.6A**). Three additional CD8α-ICL4 fusion proteins containing the S360A, S360D, or L350A/I352A substitutions were also generated.

**Figure 6.**
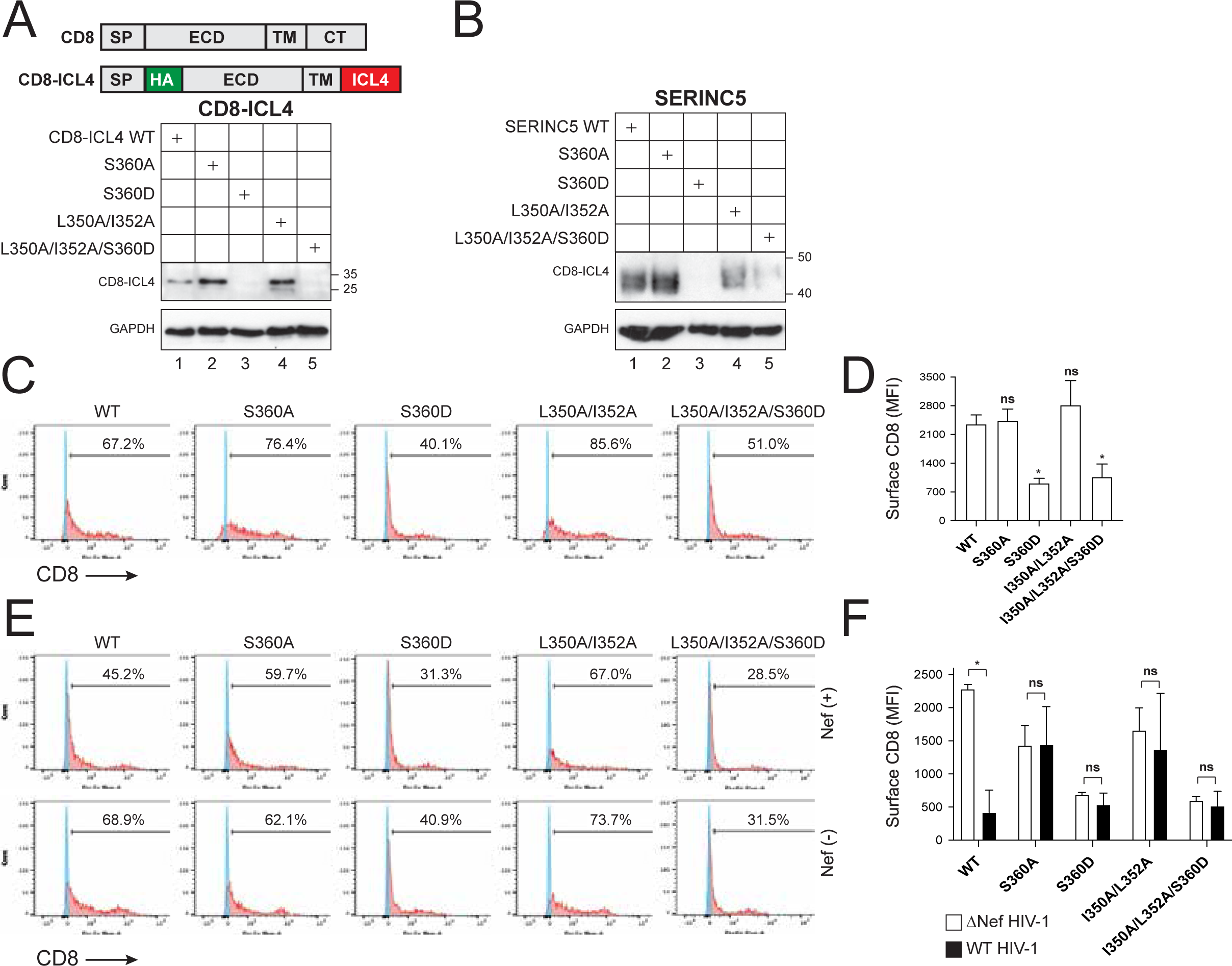
Nef downregulates CD8-ICL4 fusion protein. **A)** CD8α-ICL4 fusion proteins were created by swapping CD8α CT with ICL4 and introducing substitutions into ICL4. They were expressed in HEK293T cells, and their expression was compared by WB using anti-HA antibodies. SP, signal peptide; ECD, extracellular domain; TM, transmembrane; CT, cytoplasmic tail. **B)** Expression of SERINC5 proteins with indicated substitutions was compared by WB. **C)** Indicated CD8α-ICL4 fusion proteins were expressed in HEK293T cells and their expression on the cell surface were detected by flow cytometry after staining with fluorescent anti-HA antibodies. **D)** MFI values for the CD8-positive cell populations in C) were statistically analyzed. **E)** Indicated CD8α-ICL4 fusion proteins were expressed with WT or ΔNef HIV-1 proviral vectors in HEK293T cells and analyzed as in C). **F)** MFI values for the CD8-positive cell populations in E) were statistically analyzed. Error bars in D) and E) indicate the SEM calculated from three independent experiments. Statistical analysis: *p<0.05, ns p>0.05.

The WT, S360A, S360D, or L350A/I352A CD8α-ICL4 fusion proteins were expressed in HEK293T cells and their relative steady-state expression levels were compared. The S360A and L350A/I352A CD8α-ICL4 fusion proteins had relatively higher levels of expression compared to the WT fusion protein as compared by WB (**Fig.6A,** lane 1, 2, 4) and flow cytometry (**Fig.6C, Fig.6D**). Similar to the results described in Fig.4, for the full length S360D SERINC5, the expression of the S360D CD8α-ICL4 fusion protein was poorly detectable by WB (**Fig.6A,** lane 3). However, its low relative level of expression was confirmed by flow cytometry (**Fig.6C, Fig.6D**). These results suggest that the phosphorylation of S360 also downregulates the CD8α- ICL4 fusion protein.

To further characterize the function of the L350/I352 residues, L350A/I352A/S360D CD8α-ICL4 fusion proteins and L350A/I352A/S360D SERINC5 proteins were generated. When expressed in HEK293T cells, these triple-substitution proteins were as poorly expressed as their single S360D substitution proteins (**Fig.6B-Fig.6D**). Thus, the poor expression of the S360D substitution proteins cannot be rescued by substituting the L360/I352 residues.

Finally, we tested the ability of Nef to downregulate each of the CD8α-ICL4 fusion proteins. Nef effectively downregulated the WT, but not the S360A and L350A/I352A CD8α- ICL4 fusion proteins (**Fig.5E, Fig.5F**). The relative expression levels of the S360D and L350A/I352A/S360D CD8α-ICL4 fusion proteins were both low, regardless of Nef expression. Taken together, these results suggest that Nef downregulates CD8α-ICL4 and SERINC5 expression via a similar mechanism.

### Detection of Nef:CD8-ICL4 interaction by quantitative proteomics

To test whether Nef binds to the ICL4 of SERINC5, the WT, S360A, or L350A/I352A CD8α-ICL4 fusion proteins were co-expressed in HEK293T cells in the presence of a downregulation deficient Nef, which contains a LL/AA substitution in its dileucine motif, in order to prevent the degradation of the WT CD8α-ICL4. Next, co-immunoprecipitation (CO-IP) assays were carried out using anti-Nef antibodies. Approximately equivalent levels of each of the CD8α-ICL4 fusion proteins, as well as the Nef protein, were detected in these cells by WB (**Fig.7A**). In addition, similar levels of Nef protein were detected, as expected, whereas none of any CD8α-ICL4 fusion proteins were detected, in these IP fractions by WB (**Fig.7A**).

**Figure 7.**
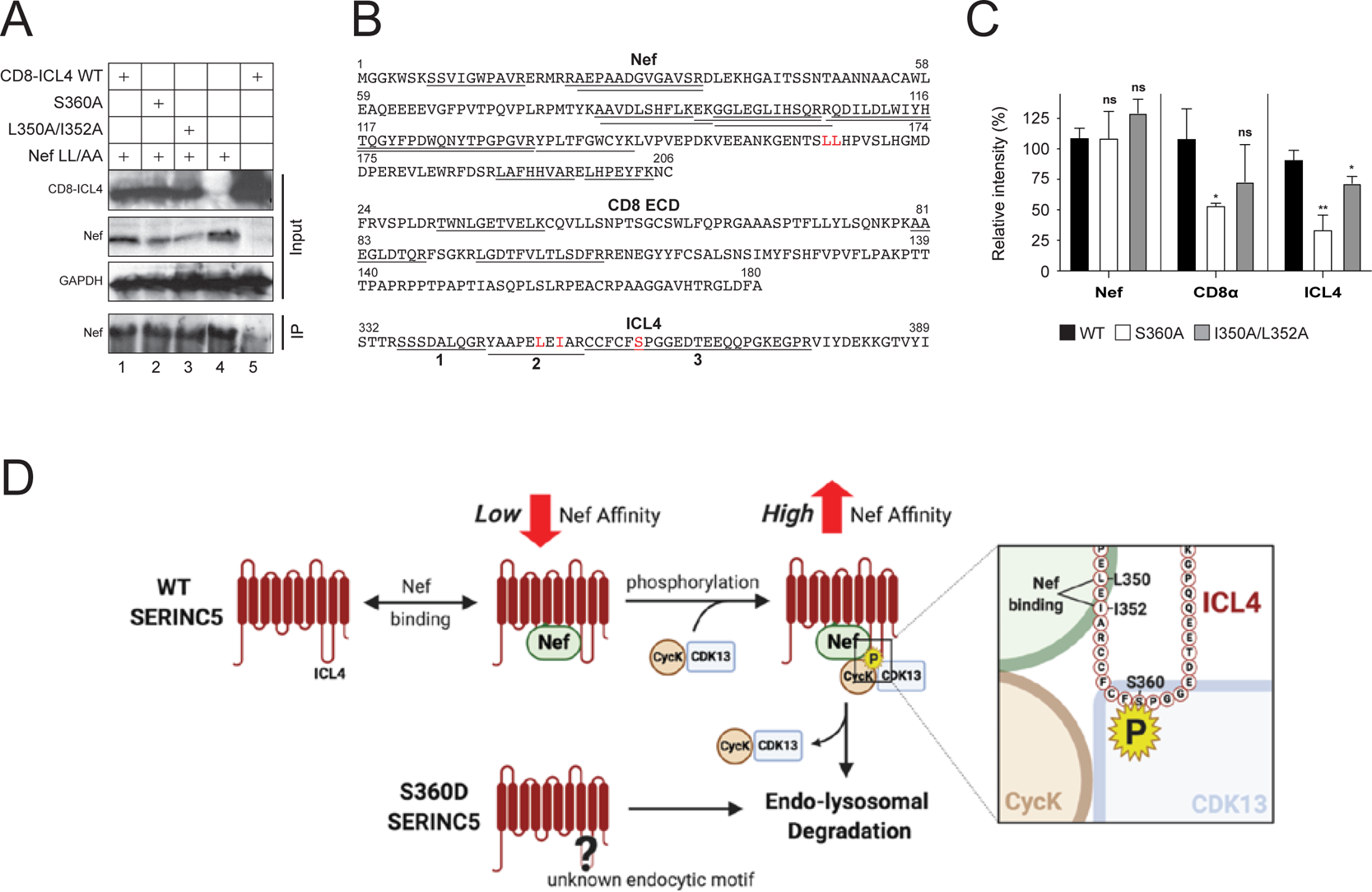
Detection of Nef:CD8-ICL4 interaction by quantitative proteomics. **A)** Indicated CD8α-ICL4 fusion proteins were expressed with the HIV-1 provirus encoding the Nef dileucine substitution (Nef LL/AA) protein. Proteins were immunoprecipitated with polyclonal anti-Nef antibodies and proteins were detected by WB using indicated antibodies. **B)** IP samples 1, 2, and 3 in A) were analyzed by mass spectrometry after trypsin digestion. Nef, CD8α, and ICL4 peptides detected from these samples are indicated in these proteins. Three ICL4 peptides are numbered (1, 2, 3). **C)** Levels of Nef, CD8α and ICL4 peptides detected by mass spectrometry in B) are quantified and presented as relative values, with the value of WT CD8α-ICL4 set to 100%. Nef proteins were quantified by label-free quantitation (LFQ) intensity. CD8α and ICL4 were quantified by the intensity of all three CD8 peptides, or ICL4 peptide-1 (SSSDALQGR), respectively. The error bars indicate the SEM calculated from three independent experiments. Statistical analysis: *p<0.05, **p<0.01, ns p>0.05. **D)** A proposed model of how S360 phosphorylation promotes SERINC5 downregulation.

We then used mass spectrometry to further analyze these IP fractions. A total of 13 Nef peptides, 3 CD8α peptides, and 3 ICL4 peptides were detected in the WT, S360A, and L350A/I352A CD8α-ICL4 derived samples (**Fig.7B**). Among the 3 ICL4 peptides, peptide-1 (^336^SSSDALQGR^344^) was detected in all three samples; peptide-2 with the L350A/I352A substitutions (^345^YAAPE<u>A</u>E<u>A</u>AR^354^) was only detectable in the L350A/I352A CD8α-ICL4 derived sample; and peptide-3 with the S360A substitution (^355^CCFCF<u>S</u>PGGEDTEEQQPGKEGPR^377^) was only detectable in the S360A CD8α-ICL4 derived sample (**Fig.S4**). Thus, we were able to detect Nef-ICL4 binding via mass spectrometry and validated the specificity of this binding by identifying each of the ICL4 substitutions in their appropriate IP fraction.

Finally, we compared the intensity of Nef binding to the WT, S360A, or L350A/I352A CD8α-ICL4 fusion proteins. We compared the label-free quantification intensity of the Nef protein, the three CD8α peptides, and ICL4 peptide-1 in each of these three IP fractions. Consistent with the WB results (**Fig.7A**), approximately equivalent quantities of Nef protein were detected in three samples, excepting a slightly higher quantity in the L350A/I352A-derived sample (**Fig.7C**). Notably, both CD8α and ICL4 quantities in the WT-derived sample were at least 2-fold higher compared to the S360A-derived sample and were also slightly higher compared to the L350A/I352A-derived sample (**Fig.7C**). These results demonstrate that S360, and to a lesser degree L350/I352, are required for Nef binding to SERINC5. Thus, we conclude that S360 phosphorylation increases SERINC5 binding to Nef.

## Discussion

In this study, we provide evidence that CycK:CDK13 is required for Nef antagonism of SERINC5. When the kinase activity of this complex was inhibited by silencing *CCNK* or *CDK13* gene expression, expressing the CDK13 kinase-inactive mutant D837A/D855N, or treating cells with the kinase inhibitor THZ531, Nef became unable to downregulate SERINC5 and/or counteract its antiviral activity. CycK:CDK13 phosphorylated SERINC5 at S360, and S360 phosphorylation was absolutely essential for these Nef activities. Thus, our findings not only reveal a new role for CycK:CDK13 in eukaryotic biology, but also extend the previous observations on interactions between Nef and CycK (Khan and Mitra, 2011). Interestingly, although CycK partners with both CDK12 and CDK13, CDK12 had no effect on SERINC5. This finding could be due to different subcellular localizations of the two CDKs or to additional interactions between CycK:CDK13, Nef, and SERINC5. Our results are consistent with those that demonstrate CDK12 and CDK13 regulate the expression of markedly different sets of genes (Greifenberg et al., 2016), We have obtained identical results from SERINC5 and the CD8-ICL4 chimeric proteins in their response to Nef, which confirms that ICL4 is targeted by Nef (Dai et al., 2018). Among those 58 aa in ICL4, S360/L350/I352 have been identified to play a critical role in Nef downregulation of SERINC5. However, because the S360D phosphomimetic SERINC5 was rapidly internalized from the cell surface and degraded in lysosomes even in the absence of Nef, ICL4 contains an unknown phosphoserine-dependent endocytic motif. This situation resembles the downregulation of CD4 from the cell surface by phorbol esters that activate protein kinase C. Activated protein kinase C phosphorylates serine residues in a dileucine motif ^408^SxIxLLS^415^ in the CD4 cytoplasmic tail, which facilitates its binding to AP-2 (Pitcher et al., 1999). As recently reported, ICL4 also binds to the μ subunit of AP-1 and AP-2 in a phosphorylation-dependent manner (Stoneham et al., 2020). We found that inactivating L350/I352 in conjunction with the S360D substitution did not rescue its expression, indicating that L350/I352 do not represent a bona fide dileucine internalization motif. Thus, in addition to S360/L350/I352, there are additional residues that play an important role in SERINC5 downregulation by interacting with adaptor proteins in a phosphoserine-dependent fashion.

We found that when L350/I352 were inactivated, the CD8-ICL4 interaction with Nef was reduced, indicating that these two hydrophobic residues are involved in SERINC5-Nef binding. Thus, although L350/I352 do not constitute a dileucine motif, they could still function as a Nef-binding site. L350/I352 should constitute a hydrophobic surface and bind to the multifunctional hydrophobic pocket of Nef, which also accommodates the CD4 dileucine motif (Kwon et al., 2020). However, because the single S360A substitution also disrupts the Nef-binding ability, the Nef- SERINC5 binding is also dependent on phosphoserine, which is different from the Nef-CD4 binding which is independent of phosphoserine (Garcia and Miller, 1991).

We suggest that S360 phosphorylation is a dynamic process, which promotes SERINC5 downregulation via changing the ICL4 conformation (**Fig.7D**). Nef interacts with SERINC5 and CycK:CDK13, which induces a conformational change in ICL4 via phosphorylating S360. This conformational change makes L350/I353 more accessible to Nef and enhances the Nef-SERINC5 binding, resulting in SERINC5 endocytosis and degradation via the Nef dileucine motif. We also suggest that this conformational change is transient due to rapid dephosphorylation by cellular phosphatases and sustainable only in the presence of Nef and CycK:CD13 (**Fig.7D**). If this conformation change is maintained, for example, by introducing the phosphomimetic S360D substitution, the unknown endocytic motif in ICL4 is stably exposed, resulting in AP2 recruitment and SERINC5 degradation in a Nef-independent manner.

In summary, our findings promise a new direction in anti-HIV therapy, namely in interfering with CDK13 as a strategy to attenuate viral replication and spread. Besides our covalent chemical inhibitor, great efforts are being made to inhibit specific transcriptional CDKs and those that function independently of the cell cycle. Our study revealed one such attractive target for HIV- 1, as CDK13 does not play an essential role in transcription or in the cell cycle.

## Acknowledgements

We thank Heinrich Gottlinger, Qintong Li, Jennifer Doudna, and the NIH AIDS Research and Reference Reagent Program for providing reagents. Graphical abstract and the model in Fig.7D were created with BioRender.com. Y.H.Z. is supported by a grant from National Institutes of Health (AI145504).

## Author Contributions

Q.C. and M.C. performed experiments for Fig.1 (B, D, E, F), Fig.2, Fig.3 (B, D, E, F), Fig.4, Fig.5 (A, B, C, D), Fig.6, Fig.7A, Fig.S1, Fig.S2, and Fig.S3. S.L., R.L., and I.A. performed experiments for Fig.3C. D.A.F performed experiments for Fig.1A. Z.Y., X.S., and L.S. performed mass spectrometry experiments for Fig.1C, Fig.5E, Fig.7 (B, C), and Fig.S4. J.H. supported purifying SERINC5 proteins. J.F.H. supported silencing *CCNK* in 293T and Jurkat cells. S.F.J. created graphical abstract and the model in Fig.7D and provided insightful comments on the manuscript. M.B.P edited manuscript. Y.H.Z. designed this study and wrote manuscript with input from all coauthors.

## Materials and Methods

### Resource Availability

#### Lead Contact

Further information and requests for resources should be directed to and will be fulfilled by the Lead Contact, Yong-Hui Zheng.

#### Materials Availability

Further information and requests for resources and reagents should be directed to Yong-Hui Zheng.

#### Data and Code Availability

Original data for figures in paper available upon request from corresponding author.

## Experimental Model and Subject Details

### Cell Lines

The human HEK293T cells were obtained from the American Type Culture Collection. TZM-bI cells were obtained from the National Institutes of Health (NIH) AIDS Research and Reference Reagent Program. HEK293T and TZM-bI cells were maintained in Dulbecco modified Eagle medium (DMEM) with 10% bovine calf serum (BCS) (HyClone), at 37°C and 5% CO2. The *SERINC3/5* knockout Jurkat-TAg (JTAg) cell line was provided by Heinrich Gottlinger and cultured in RPMI 1640 with 10% fetal bovine serum (FBS) (Sigma), at 37°C and 5% CO2.

## Method Details

### Expression vectors

The HIV-1 proviral clones pH22 and pH22ΔN were reported previously(Zheng et al., 2003). pCMV6-Ser5-FLAG, pCMV6-Ser2-FLAG, and pCMV6-CD4- FLAG were reported previously(Ahmad et al., 2019; Shi et al., 2018; Zhang et al., 2017). pBJ5- Ser5-HA and pBJ5-iHA-Ser5 were provided by Heinrich Gottlinger. pcDNA3.1-CDK12-FLAG, pcDNA3.1-CDK13-FLAG, prEF-FLAG-CCNK, pLKO.1-CCNKshRNA, pLKO.1- CDK12shRNA, pLKO.1-CDK13shRNA expression vectors were provided by Qintong Li. pCMV6-CCNK-FLAG was purchased from Origene. The Cas9 expression vector pMJ920 that has a GFP marker was obtained from Jennifer Doudna through Addgene. A *CCNK* gRNA (5’- AGGACTTGATCCAGCCACCGAGG-3’) targeting the 2^nd^ exon was expressed from pGEM-T (Promega) as we did before(Zhou et al., 2014). pcDNA3.1-CDK12-HA and pcDNA3.1-CDK13- HA were created by cloning CDK12-HA and CDK13-HA into pEGPF-N1 via HindIII/KpnI digestion. To create pGEX-ICL4, ICL4 and vector fragments were amplified from SERINC5 cDNA or pGEX-6P-1 by PCR and ligated via homologous recombination using a CloneExpress II one-step cloning kit (Vazyme Biotech, China). The S249A, S360A, S360D, T366A, and L350A/I352A mutation in pCMV6-Ser5-FLAG or pBJ5-iHA-Ser5, the L350A/I352A/S360D mutation in pBJ5-iHA-Ser5, the S360A mutation in pGEX-ICL4, the D858A and D876N mutation in pcDNA3.1-CDK12-FLAG, and the D837A and D855N mutation in pcDNA3.1-CDK13-FLAG, were created by a site-directed mutagenesis kit (Agilent). CD8α-ICL4 and its mutants were synthesized from TWIST Bioscience and expressed from the pTwist-CMV-Puro vector. All mutations were confirmed by Sanger sequencing. Primers and cloning methods are available upon request.

### Protein purification

To purify SERINC5 proteins, HEK293T cells were cultured in twenty 10-cm cell culture dishes at 2x10^6^ cells in 10 ml medium per dish. Cells were transfected with 4.5 μg pCMV6-Ser5-FLAG and 4.5 μg HIV-1 proviral vector in 27 μg PEI and cultured for 48 h. After collecting cells and washing with PBS, cells were incubated with a lysis buffer [50 mM Tris, pH 7.5, 150 mM NaCl, 5% glycerol, 1% *n*-Dodecyl β-D-maltoside (DDM, Sigma), and EDTA-free protease inhibitor cocktail (Roche)] at 4°C for 2 h. Total cell lysate was spun at 11,000 rpm using a F13-14x50cy rotor in a SORVALL RC6+ centrifuge at 4°C for 30 min. Supernatant was collected and applied to a Poly-Prep chromatography column (BioRad) containing 0.5 ml anti-FLAG M2 affinity gel (Sigma). After incubating at 4°C for 2 h with gentle shaking, supernatant was drained away, SERINC5 proteins were eluted out by 3xFLAG peptide (Sigma) at 100 μg/ml in a purification buffer (50 mM Tris, pH 7.5, 150 mM NaCl, 5% glycerol, 0.05% DDM, and protease inhibitor cocktail). SERINC5 proteins were analyzed by SDS-PAGE followed by mass spectrum and phosphorylation analyses. To purify CycK:CDK13 complex, 293T cells were transfected with an equal amount of pcDNA3.1-CDK13-FLAG and pCMV6-CCNK-FLAG. Proteins were purified similarly and used as enzymes for *in vitro* kinase assay.

To purify GST-ICL4 proteins, *E. coli* strain BL21 was transformed with pGEX-ICL4 vectors and cultured in 1 liter of LB medium at 37°C with shaking to an OD600 of 0.5-1.0. GST-fusion protein expression was induced by adding IPTG to a final concentration of 0.1 mM for an additional 3 h at 37°C with shaking. After centrifugation at 3,500 g for 20 min at 4°C, bacterial pellets were subjected to three cycles of sonication. Supernatant was collected after centrifugation of the lysate at 12,000 g for 15 m at 4°C and incubated with glutathione-Sepharose beads (GE Healthcare). Fusion proteins were eluted out by 20 mM reduced glutathione in 50 mM Tris-Cl (pH 8.0) and used as substrates for *in vitro* kinase assay.

### Immunoprecipitation (IP)

To detect protein interactions in Fig.1B and Fig.1C, FLAG-tagged proteins were expressed with their target proteins in 293T cells cultured in 10-cm dishes. Proteins were pulled down by an anti-FLAG M2 antibody (Sigma) using Pierce Classic IP Kit (Thermo Fisher Scientific) and analyzed by western blotting (WB). To detect CD8-ICL4 interactions with Nef in Fig.5D, these proteins were expressed in 293T cells cultured in 10-cm dishes. Cells were lysed in a lysis buffer [50 mM Tris, pH 7.5, 150 mM NaCl, 5% glycerol, 1% DDM, and EDTA-free protease inhibitor cocktail] at 4°C for 2 h. Cytosolic factions were collected and incubated with an anit-HIV-1 Nef polyclonal antibody (NIH AIDS Reagent Program, Cat.#2949) at 37°C for 2 h. Samples were then incubated with Protein G beads (ThermoFisher) at 4°C overnight and analyzed by WB.

### Viral antagonism of SERINC5

HEK293T cells were transfected with polyethylenimine (PEI) as we described previously(Zhang et al., 2017). Briefly, HEK293T cells were seeded in 6- well plates at 4x10^5^ cells per well with 2 ml medium one day before. Each well was transfected with 10 μg PEI mixed with 2 μg HIV-1 proviral vector and 50 ng SERINC5 expression vector in the presence or absence of 1 μg silencing vectors (Cas9+sgRNA, or shRNA vectors) or expression vectors (CycK, CDK12, CDK13). Jurkat cells were electroporated by Amaxa Nucleofector II system on program 3 for Jurkat E6 cells. Briefly, 2x10^6^ cells and 4 μg plasmid DNA were resuspended with 100 μl of Mirus electroporation buffer and electroporated. Cells were immediately transferred to 12-well plates containing 2 mL of RPMI 1640 with 10% FBS and cultured for 5 days. Relevant empty vectors were added to keep the same amounts of total DNAs for each transfection whenever necessary. Cells were collected for analyzing total protein expression by Western blotting and flow cytometry. Virions in supernatants were quantified by p24^Gag^ ELISA and viral infectivity was measured after infection of TZM-bI cells.

### Western blotting (WB)

Cells were lysed with 1% NP-40 in PBS containing protease inhibitor cocktail (Sigma). After removal of nuclei by brief centrifugation, cytosolic fraction was collected and mixed with 4x Laemmli Sample Buffer (Bio-Rad). Proteins were separated in 10% Bis-Tris acrylamide gels, blotted on to nitrocellulose membrane (Bio-Rad), and detected by antibodies. Mouse monoclonal anti-Nef was described previously(Zheng et al., 2003). Horseradish peroxidase (HRP)-conjugated mouse monoclonal anti-FLAG, anti-HA antibodies, and anti-actin were purchased from Sigma; HRP-conjugated mouse monoclonal anti-GAPDH was purchased from Proteintech; rabbit polyclonal anti-CrkRS (CDK12) and anti-CDC2L5 (CDK13) were purchased from Novus; rabbit polyclonal anti-CycK was purchased from Abcam; HRP-conjugated anti-human, -rabbit, and -mouse immunoglobulin G secondary antibodies were purchased from Thermo Fisher. Immobilon Classico and Crescendo Western HRP Substrate was purchased from MilliporeSigma.

### Viral infectivity

TZM-bI cells were seeded at 10,000 cells per well in 100 μl medium in 96-well black plates with half area transparent bottom (Greiner Bio-One). Each well was inoculated with 100 μl of viruses that were subjected to 10-fold serial dilution and infection was done in triplicate. After 48 h, viral infection was determined by a Firefly Luciferase Assay Kit (Biotium). Briefly, after removal of media, 20 μl of 1x Firefly Lysis Buffer were added to each well. After 5 m, 15 μl of Firefly Luciferase Assay Buffer 2.0 containing 0.3 μl of D-luciferin at 10 mg/ml were added to each well and luciferase activities were measured by Veritas Microplate luminometer (Turner Biosystem). Viral infectivity was calculated by normalizing luciferase activities with the amounts of p24^Gag^ in viral inoculum.

### Flow cytometry

After transfection of HEK293T and Jurkat cells with SERINC5 and other expression vectors, cells were treated with Fixation/Permeabilization Solution kit (BD Biosciences) for detecting intracellular proteins or remained untreated for detecting cell surface proteins. Cells were stained with Alexa Fluor 647- or pacific blue-labeled anti-HA antibodies (BioLegend). After extensive washing, non-permeabilized cells were fixed with 4% paraformaldehyde (PFA), and all cells were analyzed by BD LSR II.

To analyze cell cycle, Jurkat cells were transfected with increasing amounts of *CCNK* silencing or expression vectors. After 24 h, cells were collected and held in 0.4 ml 50% FBS in PBS on ice. Cells were fixed by adding 1.2 ml 70% EtOH with gentle mixing over 20-30 sec. After washing with PBS, 1 ml of Propidium Iodide Cell Cycle Reagent (Thermo Fisher) were added to stain cells for 15-30 min at 37°C and analyzed by BD LSR II. ModFit LT version 4.1.7 software was used to analyze the resulting DNA histograms and calculate the percentage of cells in each phase of the cell cycle, based on a total of 10,000 events per sample.

### Endocytosis assay

An antibody uptake assay was used to measure Nef-mediated SERINC5 endocytosis as we did previously(Shi et al., 2018). Briefly, HeLa cells were transfected with SERINC5 and Nef expression vectors and cultured for 48 h. Cells were then stained with an anti-HA antibodies to label SERINC5 proteins on the cell surface and incubated at either 37°C to allow endocytosis. One hour later, SERINC5 subcellular localization was determined by fluorescence microscopy.

### Peptides

Three peptides FCFSPGGEDTEEQQPGK, FCF{pSer}PGGEDTEEQQPGK, and FCFSPGGED<u>K</u>EEQQPGK derived from SERINC5 ICL4 were synthesized and purchased from GenScript.

### *In Vitro* kinase assay

Purified GST-ICL4 proteins or synthesized ICL4-derived peptides were pre-incubated with purified CycK:CDK13 for 10 min at room temperature in a kinase buffer [150 mM HEPES, pH 7.6, 34 mM KCl, 7 mM MgCl2, 2.5 mM dithiothreitol, 5 mM ß-glycerol phosphate, 1xPhosSTOP (Roche)] as described(Greifenberg et al., 2016). Cold ATP was added to a final concentration of 2 mM, and the reaction mixture was incubated up to 60 min at 37°C. Reactions were stopped by adding EDTA to a final concentration of 50 mM. GST-ICL4 phosphorylation was determined by phos-tag staining buffer (see below) whereas peptide phosphorylation was analyzed by capillary zone electrophoresis (CZE)-MS.

### Phos-tag gel assay

Phosphorylation of SERINC5 proteins purified from HEK293T cells in the presence of Nef, and GST-ICL4 proteins after *in vitro* kinase reaction were checked by a Phos-Tag Phosphoprotein Gel Stain kit (ABP Biosciences, Rockville, MD). Briefly, equal amounts of proteins were run on an SDS-polyacrylamide gel (PAGE) and fixed with 50% methanol and 10% acetic acid. After washing with water, the gel was incubated with staining reagent for 90 m on an orbital shaker. Background was removed by incubation with de-staining reagent and the gel was imaged using a 300-nm UV transilluminator.

### LC (liquid chromatography)-MS/MS

Proteins associated with anti-FLAG M2 affinity gels or with control beads were washed three times with PBS, and treated with 20 µl of denaturing buffer (8 M urea, 100 mM NH4HCO3) at 37 °C for 30 min. Proteins were then reduced with addition of 2 µl of 5 mM dithiothreitol (DTT) for 20 m at 37 °C. After alkylation with addition of 5 µl of 5 mM iodoacetamide (IAA) for 10 min at room temperature in dark, the sample was diluted 4 times with addition of 60 µL of 100 mM NH4HCO3 to reduce the concentration of urea. After digestion with 0.2 µg of trypsin at 37 °C for overnight, 1 µl of formic acid (FA) was added to quench the digestion. The sample tube was centrifuged at 1000 rpm for 30 seconds and supernatant was taken out for desalting using Zip-tip (Millipore, ZTC18S096). Protein digest was resuspended in 10 µl of buffer A [2% Acetonitrile (ACN), 0.1% FA] and 1 µl of sample was loaded for nanoRPLC-MS/MS analysis.

An EASY nanoLC-1200 system (Thermo Fisher) equipped with a C18 RPLC column (75 µm i.d. x 50 cm, C18, 2 µm, 100 Å, Thermo Fisher Scientific) was connected to a Q-Exactive HF mass spectrometer (Thermo Fisher) for nanoRPLC-MS/MS analysis. Buffer A containing 2% ACN and 0.1% FA, and buffer B containing 80% (v/v) ACN and 0.1% FA were used to generate gradient separation. The flow rate was 200 nl/min. The gradient for RPLC separation was as follows: from 8 to 30% (v/v) B in 50 min, from 30% to 50% (v/v) B in 30 min, from 50% to 80% (v/v) B in 10 min and maintain at 80% (v/v) B for 10 min. A Top10 data dependent acquisition (DDA) method was used. The MS parameter was set as follows: the full MS resolution was 60,000 and the maximum injection time was 50 ms. The scan range was 300 – 1800 m/z. The AGC target was 3e6. The MS/MS resolution was 30,000 and the maximum injection time was 50 ms. The AGC target was set 1e5 for MS/MS. The isolation window was set 2.0 m/z. The intensity threshold for fragmentation was 5e4. The dynamic exclusion was set 30 s.

### Database Search for protein identification

Proteome Discoverer 2.2 (Thermo Fisher) with SEQUEST HT searching engine built in was used. The mass tolerance for precursor ion was set 20 ppm and for fragment ion was set 0.05 Da. UniProt human proteome database (UP000005640) was used for the database searching. Trypsin was set as the enzyme with two maximum missed cleavages. Oxidation on Methionine, acetylation on Protein N-terminal, phosphorylation on Serine, Tyrosine and Threonine and deamination on Asparagine or Glutamine were set as variable modifications. Carbamidomethylation on cysteine was set as the fixed modification. The peptides were filtered with confidence as high, corresponding to a 1% peptide-level false discovery rate. Protein grouping was enabled, and the strict parsimony principle was applied.

### Capillary zone electrophoresis (CZE)-MS

The synthesized peptide samples from the CCNK/CDK13 incubation and mock treatment experiments were desalted using C18 ZipTips (MilliporeSigma), lyophilized, and dissolved in 50 mM ammonium bicarbonate. A commercialized electro-kinetically pumped sheath flow CE-MS interface (an EMASS-II CE-MS interface, CMP Scientific) was employed to couple CZE with MS(Sun et al., 2015). A 7100 CE System (Agilent) was used for the automated operation of CZE. The electrospray (ESI) emitters were pulled from borosilicate glass capillary (1.0 mm o.d., 0.75 mm i.d.) by a Sutter P-1000 flaming/brown micropipette puller. The opening of the emitter was 20-40 µm. Voltage for ESI was ∼2.2 kV.

A 1-meter-long fused silica capillary (50 µm i.d., 360 µm o.d.) was used for CZE separation. The inner wall of the capillary was coated with linear polyacrylamide (LPA) as reported(Chen et al., 2017). The background electrolyte (BGE) for CZE was 5% (v/v) acetic acid (pH 2.4) and the sheath buffer for electrospray was 10% (v/v) methanol and 0.2% (v/v) formic acid in water. For each run, about 500-nl sample was injected into the capillary. Then 30 kV was applied at the injection end and 100 mbar was applied at the mean time for CZE separation for 35 min.

A 6545XT AdvanceBio Q-TOF mass spectrometer (Agilent) was used for data acquisition. The gas temperature was 325 °C. The drying gas was 1 l/min. The fragmentor and skimmer were set as 150 V and 65 V, respectively. The mass range was 800-1000 *m/z*. The acquisition rate was 0.4 spectra/s. The acquisition mode was Extended Dynamic Range (2 GHz) and the slicer mode was High sensitivity. The Agilent MassHunter Qualitative Analysis Navigator B.08.00 was used for data analysis.

### Statisitical Analysis

Statistical tests were performed using GraphPad Prism 8. Variance was estimated by calculating the standard deviation (SD) andrepresented by error bars. Significance of differences between samples was assessed using two-way ANOVA with Bonferroni post-test. All experiments were performed independently at least three times, with representative experiment being shown. *p<0.05, **p<0.01, ***p<0.001, ns, not significant (p>0.05).

## Supplemental Figures

**Figure S1.**
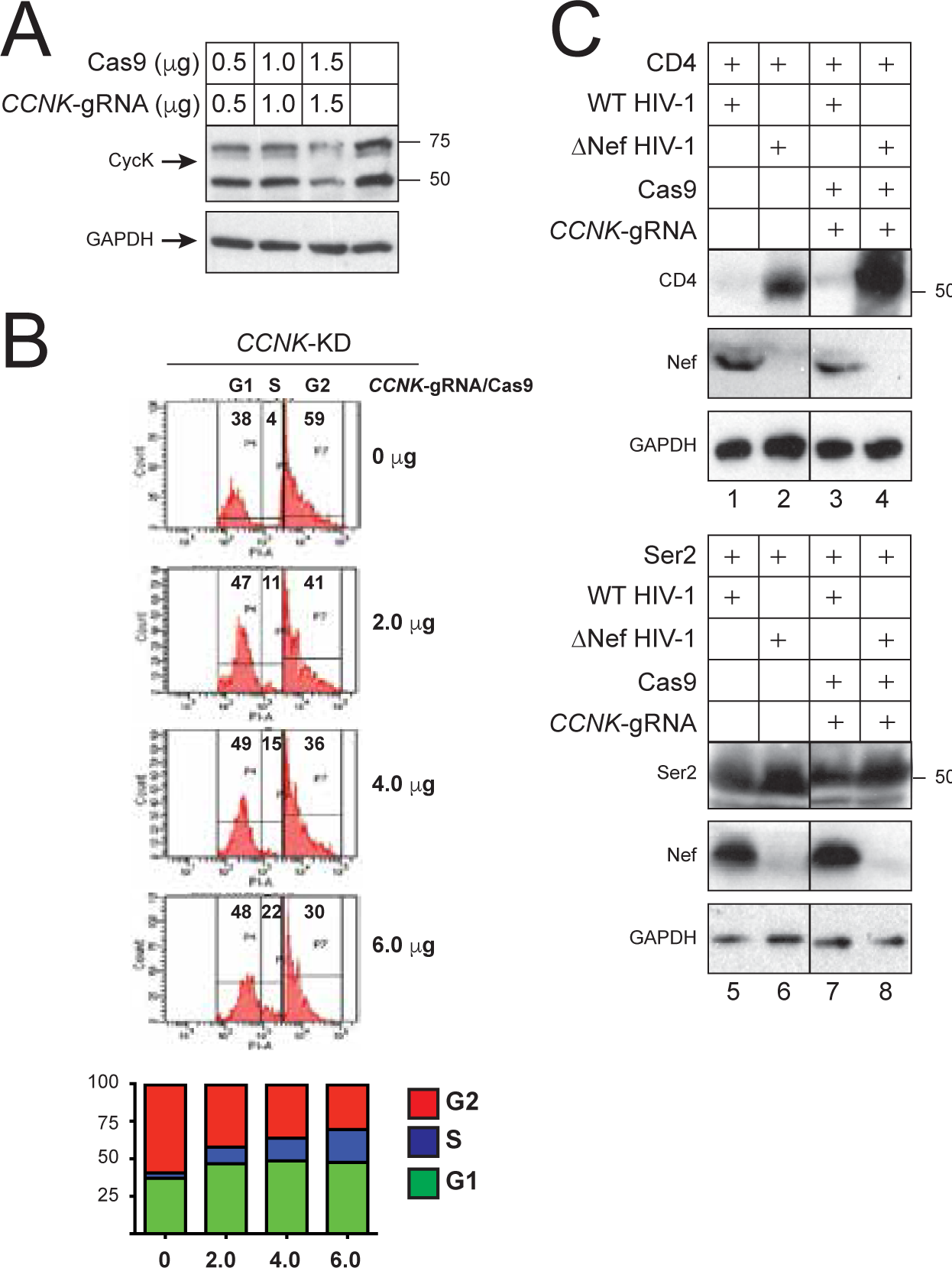
Effectiveness of CycK knockdown by CRISPRi/Cas9. **A)** HEK293T cells were transfected with increasing amounts of Cas9 and *CCNK*-gRNA expression vectors. CycK expression was detected by WB using anti-CycK antibodies. **B)** *SERINC3/5* knockout Jurkat-TAg cells were transfected with increasing amounts of Cas9/*CCNK*-gRNA expression vectors. After 48 h, cell cycle was analyzed by flow cytometry. Representative individual cell cycle distributions are shown in the histogram. **C)** CD4 and SERINC2 were co-expressed with WT and ΔNef HIV-1 proviral vectors in the presence of Cas9/*CCNK*-gRNA expression vectors in HEK293T cells. CD4, SERINC2, and CycK were detected by WB using anti-FLAG antibodies.

**Figure S2.**
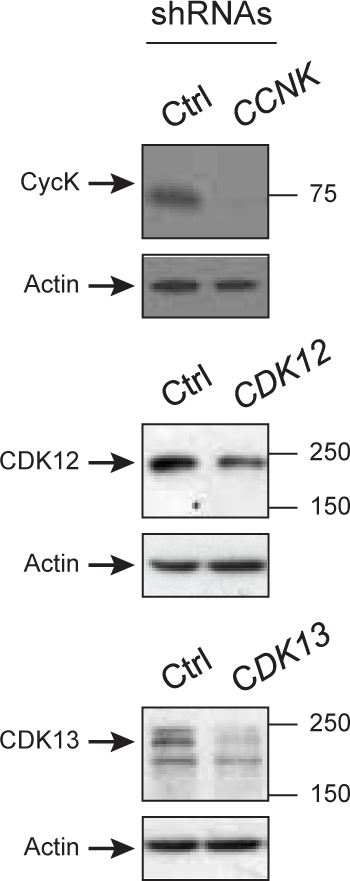
Effectiveness of shRNAs on CycK, CDK12 and CDK13 knockdown. HEK293T cells were transfected with shRNA expression vectors against *CCNK*, *CDK12* or *CDK13.* A scrambled shRNA expression vector was used as the control. CycK, CDK12 and CDK13 expression were detected by WB using their specific antibodies.

**Figure S3.**
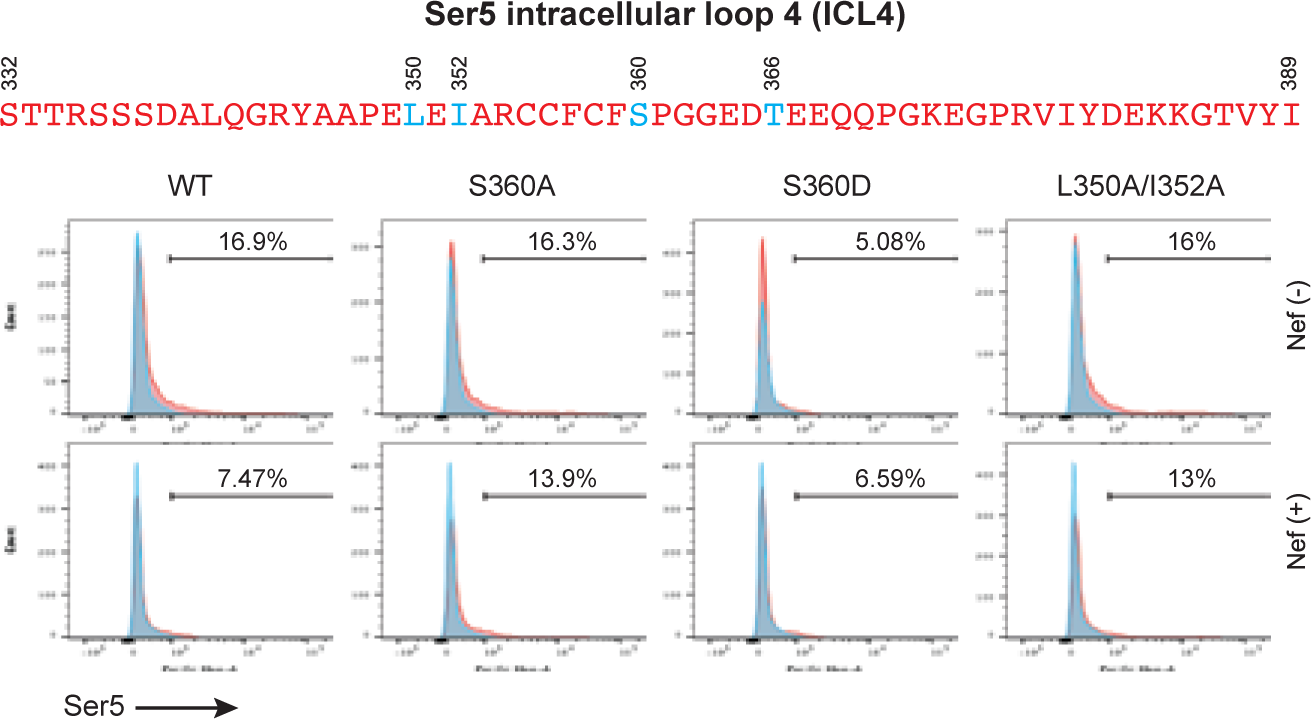
S360 is required for Nef downregulation of SERINC5 in HEK293T cells. WT SERINC5 and mutant SERINC5 S360A, S360D, and L350A/I352A proteins with an internal HA-tag were expressed alone, or with WT or ΔNef HIV-1 proviruses in HEK293T cells. Cells were stained with fluorescent anti-HA antibodies, and SERINC5 expression was determined by flow cytometry.

**Figure S4.**
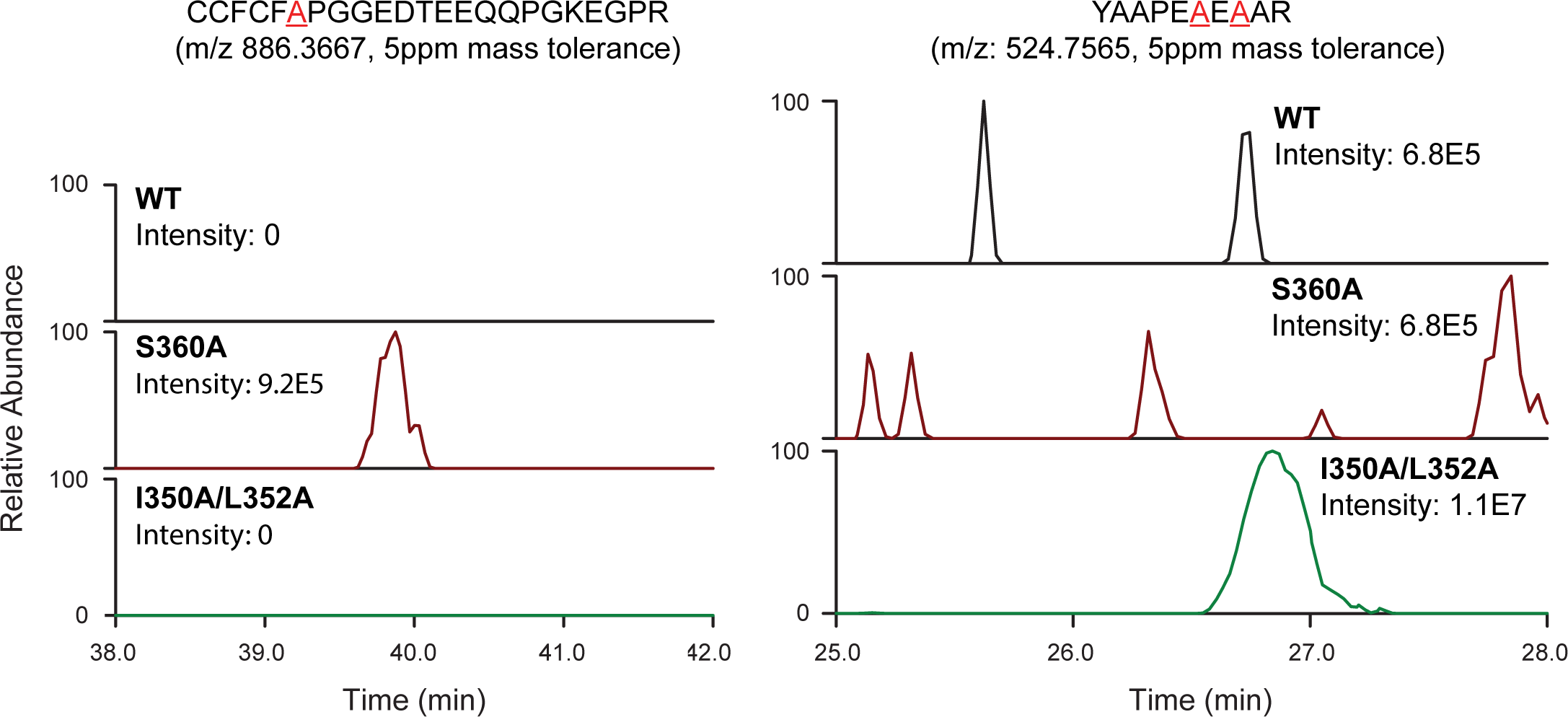
Extracted ion chromatogram (EIC) of two ICL4 peptides containing S360A or I350A/L352A mutations. EICs of peptide CCFCF<u>A</u>PGGEDTEEQQPGKEGPR and YAAPE<u>A</u>E<u>A</u>AR detected in samples from WT CD8α-ICL4 fusion protein, as well as mutant CD8α-ICL4 S360A, and I350A/L352A chimeras in are presented. The S360A and I350A/L352A mutations are underlined and shown in red.

